# LM11a-31 Inhibits p75 Neurotrophin Receptor (p75^NTR^) Cleavage and is Neuroprotective in a Cell Culture Model of Parkinson’s Disease

**DOI:** 10.1101/2024.09.10.612299

**Authors:** Poshan V. Pokharel, Aaron M. Newchurch, Sunny C. Overby, Cassidy A. Spease, Lorelei G. Darzi, Bradley R. Kraemer

**Affiliations:** Department of Biological Sciences, University of Alabama in Huntsville, Huntsville, AL 35899; Department of Biological Sciences, Eastern Kentucky University, Richmond, KY 40475

**Author notes:** **Correspondence:** Bradley R. Kraemer; Address: 369P Shelby Center for Science and Technology, 301 Sparkman Dr., Huntsville, AL 35899.

## Abstract

The p75 Neurotrophin Receptor (p75^NTR^) is a multifunctional transmembrane protein that mediates neuronal responses to pathological conditions in specific regions of the nervous system. In many biological contexts, p75^NTR^ signaling is initiated through sequential cleavage of the receptor by α- and γ-secretases, which releases receptor fragments for downstream signaling. Our previous work demonstrated that proteolytic processing of p75^NTR^ in this manner is stimulated by oxidative stress in Lund Human Mesencephalic (LUHMES) cells, a dopaminergic neuronal cell line derived from human mesencephalic tissue. Considering the vulnerability of dopaminergic neurons in the ventral mesencephalon to oxidative stress and neurodegeneration associated with Parkinson’s disease (PD), we investigated the role of this signaling cascade in neurodegeneration and explored cellular processes that govern oxidative stress-induced p75^NTR^ signaling. In the present study, we provide evidence that oxidative stress induces cleavage of p75^NTR^ by promoting c-Jun N-terminal Kinase (JNK)-dependent internalization of p75^NTR^ from the cell surface. This activation of p75^NTR^ signaling is counteracted by tropomyosin-related kinase (Trk) receptor signaling; however, oxidative stress leads to Trk receptor downregulation, thereby enhancing p75^NTR^ processing. Importantly, we demonstrate that this pathway can be inhibited by LM11a-31, a small molecule modulator of p75^NTR^, thereby conferring protection against neurodegeneration. Treatment with LM11a-31 significantly reduced p75^NTR^ cleavage and neuronal death associated with oxidative stress. These findings reveal novel mechanisms underlying activation of p75^NTR^ in response to oxidative stress, underscore a key role for p75^NTR^ in dopaminergic neurodegeneration, and highlight p75^NTR^ as a potential therapeutic target for reducing neurodegeneration in PD.

## Introduction

Neurotrophins are structurally related, diffusible proteins that regulate a variety of neuronal functions, including differentiation, neurite outgrowth, synaptic plasticity and survival[1]. The four members of the neurotrophin family include Nerve Growth Factor (NGF), brain-derived neurotrophic factor (BDNF), neurotrophin-3 (NT3), and neurotrophin-4 (NT4). These proteins exert their effects through activation of two types of receptors: tropomyosin-related kinase (Trk) receptors, including TrkA, TrkB, and TrkC, or the p75 Neurotrophin Receptor (p75^NTR^). Trk receptors function as receptor tyrosine kinases, and their activation by neurotrophins induces their trans-autophosphorylation, leading to downstream cascades that promote neuronal survival, neurite outgrowth, or synaptic strengthening, among other functions[2]. The physiological roles of p75^NTR^ are also diverse, and, with exceptions, often opposite to those of Trk receptors. For example, p75^NTR^ has been demonstrated to inhibit neurite outgrowth[3, 4], promote long-term depression[5, 6], and induce neuronal death[7-9]. Further contributing to their functional diversity, neurotrophins are initially synthesized as larger precursors, known as proneurotrophins, which can be cleaved in the Golgi apparatus or extracellular milieu to yield mature proteins. Under certain pathological conditions, uncleaved proneurotrophins, which have minimal affinity for Trk receptors, can still stimulate p75^NTR^ signaling by engaging with a complex that includes p75^NTR^ and its coreceptor, sortilin[10-12].

While signaling cascades activated by Trk receptors have been well-characterized[13, 14], the interactions through which p75^NTR^ exerts its effects are more poorly understood. This is in part due to the receptor’s complex array of interactors with expression profiles that vary by cell type[15, 16]. However, one well-established p75^NTR^ signaling mechanism that has been identified across a variety of cell types involves regulated intramembrane proteolysis of the receptor. Within this process, the ectodomain of p75^NTR^ is cleaved by an enzyme in the A Disintegrin and Metalloproteinase (ADAM) family, yielding a membrane-bound C-terminal fragment (p75^NTR^-CTF). The p75^NTR^-CTF is susceptible to cleavage by the γ-secretase complex, which releases the intracellular domain of p75^NTR^ (p75^NTR^-ICD) for interaction with various cytosolic interactors.

Proteolytic processing of p75^NTR^ in this manner has been linked to a variety of functional outcomes, including promotion of neuronal death in sympathetic and hippocampal neurons[17, 18], inhibition of neurite outgrowth in cerebellar neurons[19], proliferation of glioma cells[20], and regulation of angiogenesis after retinal hypoxia[21]. However, the effects of coreceptors, such as Trk receptors and sortilin, on p75^NTR^ processing remain poorly understood.

Among its many roles, p75^NTR^ serves a key mediator of neuronal death associated with pathological conditions. For example, the receptor has been reported to induce apoptosis of hippocampal neurons in response to seizures[17], oligodendrocytes after spinal cord injury[22], and cholinergic neurons after cortical brain injury[23]. Moreover, p75^NTR^ signaling has been implicated in neurodegeneration in models of Alzheimer’s disease[24, 25], Huntington’s disease[26, 27], and amyotrophic lateral sclerosis[28-30]. This accumulation of evidence has fostered interest in p75^NTR^ as a potential pharmacological target for mitigating neurodegeneration[31, 32]. However, the effects of the receptor on neurodegeneration appear to be context- and cell type-dependent. For example, rather than inducing death, the receptor can stimulate pro-survival signaling cascades in developing cerebellar granular neurons and trigeminal neurons[33, 34]. Thus, understanding the impact of p75^NTR^ on a particular neurodegenerative condition requires characterizing the effects of the receptor in the cell populations most vulnerable to the disease.

Previous reports have indicated that p75^NTR^ is produced in dopaminergic neurons of the ventral mesencephalon, a cell population vulnerable to oxidative stress and neurodegeneration associated with Parkinson’s disease (PD)[35-38]. Despite these findings, the role of p75^NTR^ in neurodegeneration associated with PD remains unclear. Our earlier research demonstrated that p75^NTR^ processing is induced by oxidative stress in a cell culture model of PD[39]. However, the specific signaling events that govern oxidative stress-induced p75^NTR^ signaling are not well-understood, and the physiological significance of p75^NTR^ cleavage in dopaminergic cells remains undetermined. In the present study, we reveal receptor internalization and downregulation of Trk signaling as key processes facilitating p75^NTR^ cleavage induced by oxidative stress in dopaminergic cells. With further analyses, we show that this pathway can be inhibited by LM11a-31, a small molecule modulator of p75^NTR^. Importantly, we demonstrate that treatment with LM11a-31 protects dopaminergic cells from neurodegeneration associated with oxidative stress, thus revealing a pro-death role for p75^NTR^ processing in dopaminergic cells and highlighting p75^NTR^ as a potential pharmacological target for reducing dopaminergic neurodegeneration.

## Methods

### Cell Culture

Lund human mesencephalic (LUHMES) cells (ATCC, Manassas, VA, USA, RRID: CVCL_B056) were cultured in round plastic cell culture dishes (USA Scientific, Ocala, FL, USA) or Lab-Tek II 8-well chambered slides (Thermo Fisher Scientific, Waltham, MA, USA), as previously described[39, 40]. Prior to plating, all culture containers were coated with 100 μg/mL poly-L-ornithine (Sigma-Aldrich, St. Louis, MO, USA, Cat no: P3655) at room temperature overnight and then incubated with 2 μg/mL fibronectin (Sigma-Aldrich, Cat no: F0895) for 3 hours at 37 °C. Glass chamber slides were additionally coated with 100 ng/mL poly-D-lysine (Sigma-Aldrich, Cat no: P7280) and 10 μg/mL laminin (Corning, NY, USA Cat no: 354232) at room temperature overnight to promote cell adhesion. Cells were initially seeded in growth medium consisting of Dulbecco’s Modified Eagle Medium with Nutrient Mixture F-12 (DMEM/F12) (Gibco, Waltham, MA, USA Cat no: 11330057), supplemented with 2 mM glutamine (VWR, Radnor, PA, Cat no: VWRL0131-0100), 1% (v/v) N-2 supplement (Gibco, Cat no: 17-502-048), and 40 ng/mL basic fibroblast growth factor (bFGF) (R&D Systems, Minneapolis, MN, USA, Cat no: 3718-FB). To differentiate the cells into post-mitotic neurons, the cultures were reseeded in differentiation medium, consisting of DMEM/F12 containing 2mM glutamine (VWR), 1% (v/v) N-2 supplement, 1 mM N6, 2’-O-Dibutyryladenosine 3’,5’-cyclic monophosphate (db-cAMP) (Enzo Life Sciences, Farmingdale, NY, USA Cat no: BML-CN125-0100), 2 ng/mL glial cell line-derived neurotrophic factor (GDNF) (R&D Systems, Minneapolis, MN, USA, Cat no: 212-GD-010), and 1 μg/mL tetracycline (Sigma-Aldrich, St. Louis, MO, USA, no: 87128-25G). Cultures were maintained at 37 °C and 5% CO2, and a half-volume media change was performed every other day until treatment. Cell stocks with passage numbers of 6 or less were used for all experiments to minimize genetic drift.

### Cell Treatment

After 5 days of differentiation, LUHMES cells were subjected to a full media change and then treated with vehicle solution or 6-hydroxydopamine (6-OHDA) (Sigma-Aldrich, Cat no: 162957) at the indicated concentrations. 6-OHDA solution was prepared in cold, phosphate-buffered saline (PBS)(Corning, Manassas, VA, USA, Cat no: 45000-448) with 0.02% ascorbate (Sigma-Aldrich, St. Louis, MO, USA, Cat no: A5960) to prevent autoxidation. For experiments involving JNK inhibition, cultures were pretreated with 10 µM SP600125 (Sigma-Aldrich, St. Louis, MO, USA, Cat no: A5960) for one hour prior to 6-OHDA exposure. During experiments evaluating the effects of receptor internalization of p75^NTR^ processing, cultures were cotreated with 80 µM dynasore (Sigma-Aldrich, St. Louis, MO, USA, Cat no: D7693) or vehicle solution consisting of dimethyl sulfoxide (DMSO, VWR, Radnor, PA, USA, Cat No. 97063-136). In assays involving inhibition of Trk signaling, the cultures were treated with 5 µM LOXO-101(Cayman Chemical Company, Ann Arbor, MI, USA, Cat No. 27056), a Trk receptor inhibitor, or the vehicle solution DMSO, for 3 hours prior to exposure to 10 µM 6-OHDA, and LOXO-101 was retained in the media during 6-OHDA exposure. For experiments involving small molecule modulation of p75^NTR^, cultures were cotreated with 20 nM (2S,3S)-2-Amino-3-methyl-N-[2-(4-morpholinyl)ethyl]pentanamide dihydrochloride (LM11a-31)(R&D Systems, Minneapolis, MN, USA, Cat no: 5046) or vehicle solution consisting of DMSO during application of 6-OHDA. All drugs were stored at -20o C under inert gas and protected from light.

### Immunoblotting Assays

LUHMES cell cultures were established in plastic culture dishes, differentiated for five days, and treated for 18 hours as described. Cultures were then lysed in buffer consisting of 25 mM Tris (pH 7.4),137 mM NaCl, 2.7 mM KCl, 1% Igepal CA-630, and 10% glycerol supplemented with a Complete Mini EDTA-free protease inhibitor mixture tablet (Roche, Basel, Switzerland, Cat No: 11836170001) and a PhosStop phosphatase inhibitor mixture tablet (Roche, Cat no: 4906837001). Total protein content was measured by Bradford assay using Coomassie blue reagent (Bio-Rad Laboratories Inc., Hercules, CA, Cat no. 5000006). Samples containing 40 µg protein were dissolved in SDS sample buffer (final concentration of 62.6 mM Tris-HCl, 2% sodium dodecyl sulfate, 10% glycerol, 0.005% bromophenol blue, and 1.54% dithiothreitol) and denatured by heating for 5 minutes at 95o C. Samples were then subjected to SDS-PAGE and transferred to PVDF membrane. Blotting was performed using primary antibodies for p75NTR-ICD (a generous gift from Dr. Bruce Carter, Vanderbilt University), sortilin (R&D Systems, Cat No. MAB3154), TrkA (Cell Signaling Technology, Danvers, MA, USA, Cat No. 2505), TrkB (Cell Signaling Technology, Cat No.

4603), TrkC (Cell Signaling Technology, Cat No. 3376), actin (Sigma-Aldrich, MAB1501, RRID:AB_2223041) or tubulin (Covance, Princeton, NJ; RRID:AB_2313773) and secondary antibodies conjugated to horseradish peroxidase (Jackson ImmunoResearch Laboratories, West Grove, PA, USA, Cat No. 111-035-144 and 115-035-146, RRID:AB_2307391 and AB_2307392). Blots were visualized using a Chemidoc MP gel documentation system (Biorad Laboratories Inc., Hercules, CA, USA). Densitometric analyses were performed using Image Lab Software (Bio-Rad Laboratories Inc., Version 6.0.0), and intensity measurements were normalized to actin or tubulin values as indicated.

### Immunostaining

LUHMES cells were cultured in Lab-tek II 8-well chamber slides, differentiated for 5 days, and treated as indicated. Fixation of cells with 4% paraformaldehyde in PBS was performed 24H after treatment. Fixed slides were blocked with 10% normal goat serum in PBS and then stained with Wheat Germ Agglutinin (WGA) – CF-568 conjugate (Biotium, San Francisco, CA, USA, Cat No.

29077-1), primary antibody for p75^NTR^-ECD (Cell Signaling Technology, Cat No. 8238), secondary antibody conjugated to Alexa Fluor 488 (Thermo Fisher Scientific; 1:1000. RRID: AB_142495), and/or 4’, 6’-diamidino-2-phenylindole (DAPI, 5 µg/mL, Sigma Aldrich, Cat No. D9542). For cell survival assays, permeabilization of the cells was performed during staining by including 0.1% Triton-X-100 (Sigma Aldrich, Cat No. T9284) in the blocking solution, primary antibody solution, and wash buffer. For analysis of receptor internalization, staining of unpermeabilized cells was performed through exclusion of Triton-X-100 from all staining solutions. Glass coverslips were applied to all slides using Fluoromount-G mounting solution (Southern Biotech, Birmingham, AL, USA) before visualization by confocal microscopy.

### Receptor Internalization Analysis

Unpermeabilized cells were stained for WGA and p75^NTR^-ECD as described, followed by visualization with a Zeiss LSM 700 confocal microscope system. Using Zen 2 software (Zeiss, Oberkochen, Germany), confocal micrographs were captured at a z-depth near the center region of Different cells, enabling visualization of the cell surface with the WGA stain. Using Image J software with the Fiji package, outlines of the WGA-labeled cell surface of each cell were manually traced, and mean fluorescence intensity within the outlined regions was then calculated for the channel corresponding to p75^NTR^-ECD labeling.

### Cell Viability Analysis

After the indicated treatments, LUHMES cells cultured in 8-well chamber slides were assessed for viability as previously described[39, 41]. Briefly, slides were fixed with 4% PFA, and DAPI staining was performed to label cell nuclei. The 20X objective of an LSM700 confocal microscope system (Zeiss, Oberkochen, Germany) was used to capture micrographs of cell nuclei. While healthy cells have large nuclei with lower staining intensity, dying cells exhibit nuclear envelope collapse, chromatin condensation, and nuclear fragmentation, resulting in a nuclear stain that appears brighter, smaller, and fragmented. Based on these morphological features, nuclei were scored as healthy or dying by an investigator blinded to the treatment conditions. Counts were obtained from at least five fields of view per well, with at least 125 total cells counted per condition in each of the six independent experiments.

### Statistical Analyses

Statistical analyses and graphing of data were performed using Graphpad Prism 10.3.0. Quantitative data are represented as box and whisker plots, with boxes encompassing the 25^th^ – 75^th^ percentiles, whiskers reflecting the 10^th^ – 90^th^ percentiles, and a middle line representing the median. All data were tested for normality by Shapiro-Wilk test. For normal datasets, comparison of means between groups was performed by one-way analysis of variance (ANOVA) with a Tukey’s Honestly Significant Difference (HSD) post-hoc analysis. For data with non-normal distribution, comparisons were made by a Friedman test with a Dunn’s test for multiple comparisons. P values are reported throughout the text, and statistical significance was accepted at P < 0.05.

### Ethical Statements

Institutional ethical approval of the experiments was not required, as the present study did not involve the use of animals or human subjects. The authenticity of the LUHMES cells was confirmed by assessing their ability to differentiate into post-mitotic neurons, as well as by confirming their expression of ßIII-tubulin and tyrosine hydroxylase after five days of differentiation.

## Results

### Oxidative Stress Stimulates JNK-Dependent Endocytosis of p75^NTR^

We previously discovered that oxidative stress triggers proteolytic processing of p75^NTR^ in a cell culture model of Parkinson’s disease (PD). Notably, this signaling process depends on c-Jun N-terminal Kinase (JNK), a stress-activated kinase[39]. Since JNK has been shown to facilitate the endocytosis of p75^NTR^ in sympathetic neurons treated with BDNF[42], we predicted that oxidative stress would similarly induce JNK-dependent internalization of p75^NTR^ in our PD cell culture model. To test this, we used immunofluorescence and confocal microscopy to examine the presence of p75^NTR^ on the cell surface of differentiated LUHMES cells subjected to oxidative stress with or without prior JNK inhibition. Compared to healthy cells, a significant decrease (P = 0.0269) in p75^NTR^ at the cell surface was observed in cells treated with 6-hydroxydopamine (6-OHDA) to induce oxidative stress. In contrast, cells pretreated with the JNK inhibitor SP600125 before 6-OHDA exposure exhibited significantly higher (P = 0.0380) levels of cell surface-localized p75^NTR^ (Fig. 1A -1B). These findings indicate that oxidative stress promotes internalization of p75^NTR^ through a mechanism requiring JNK.

**Figure 1.**
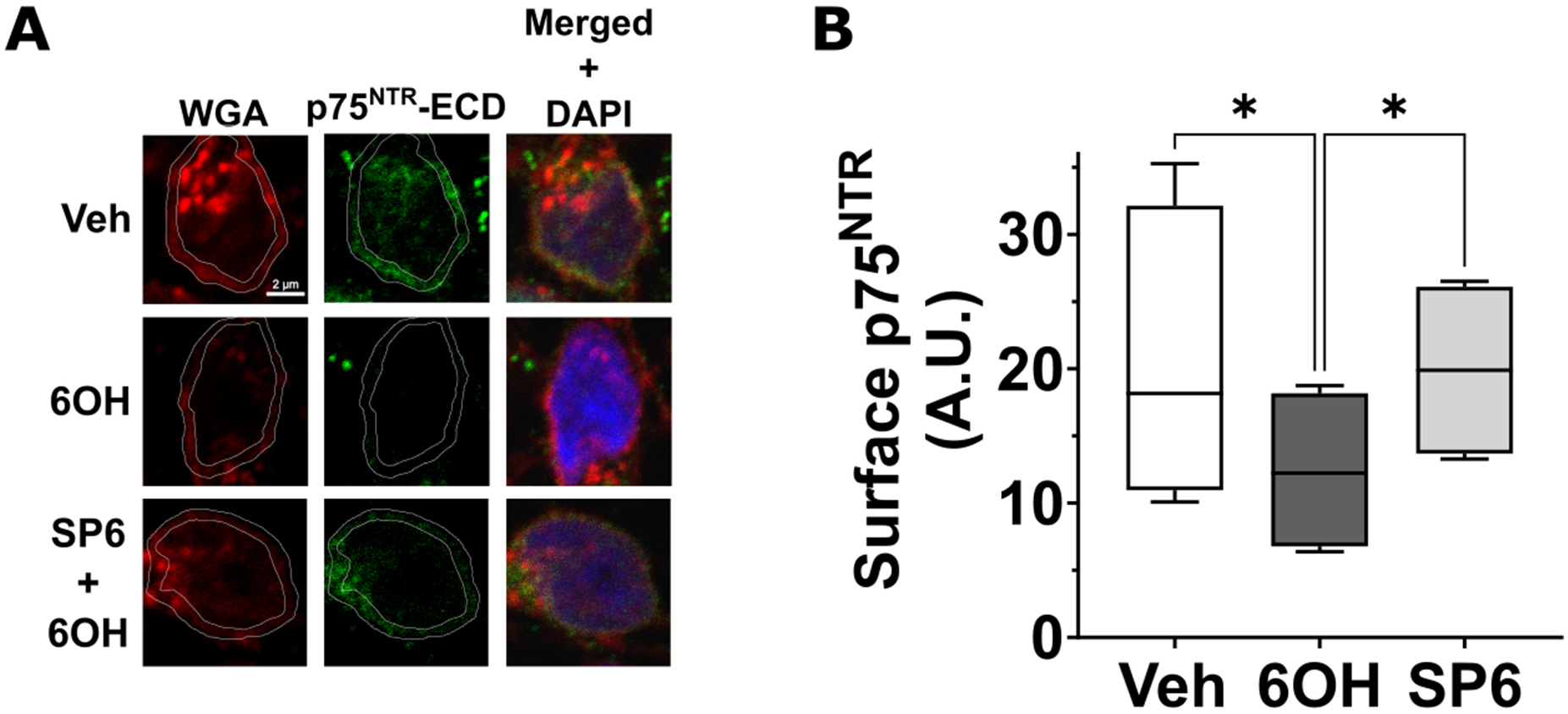
6-OHDA Induces JNK-mediated Endocytosis of p75^NTR^. **(A)** Confocal micrographs of cell surface p75^NTR^ (green) and plasma membrane (red) staining in LUHMES cells that were differentiated for five days and then treated for 18H with 7.5 µM 6-OHDA or vehicle solution following a one-hour pretreatment with 10 µM SP600125 or vehicle solution. Cell surface p75^NTR^ was labeled by immunostaining fixed, unpermeabilized cells with an antibody specific for the p75^NTR^-ECD. Plasma membrane labeling was performed through staining with Wheat germ agglutinin-CF®568 conjugate, and nuclei (blue) were labeled with 4’,6-diamidino-2-phenylindole (DAPI). **(B)** Quantification of cell surface-localized p75^NTR^ in LUHMES cells that were treated, stained, and imaged as described in 1A (*n* = 4, ANOVA with Tukey’s HSD). Abbreviations: *Veh*, vehicle; *6OH*, 6-hydroxydopamine; *SP6*, SP600125; *A.U*., arbitrary units; *, p<0.05; *n*, number of experiments, each featuring an independent cell culture preparation.

### Oxidative Stress-Induced p75^NTR^ Processing Requires Receptor Endocytosis

Based on our observation that oxidative stress promotes endocytosis of p75^NTR^, we next considered whether this internalization of the receptor influences its proteolytic processing.

Differentiated LUHMES cells were subjected to oxidative stress by exposure to 6-OHDA, and the cells were cotreated with vehicle solution or dynasore, a dynamin inhibitor that has previously been demonstrated to block endocytosis of p75^NTR^ [42-46]. Lysates were assessed for p75^NTR^ cleavage by western blot using an antibody specific for the intracellular domain of p75^NTR^ (Fig. 2A and 2B). In alignment with our previous findings, cleavage of p75^NTR^ was observed in cells subjected to oxidative stress, as our densitometric analyses revealed a significant accumulation (P = 0.0099) of p75^NTR^-C-terminal fragment (p75^NTR^-CTF) and p75^NTR^-intracellular domain (p75^NTR^-ICD) (P = 0.0028), as well as a significant decrease (P = 0.0273) in full-length p75^NTR^, in cells exposed to 6-OHDA compared to healthy cells (Fig. 2C – 2E). However, in cells co-treated with dynasore, proteolytic processing of p75^NTR^ was reduced, as indicated by a significant reduction in p75^NTR^-CTF (P = 0.0226) and p75^NTR^-ICD (P = 0.0083) (Fig. 2C and 2D). Moreover, this reduction in p75^NTR^ cleavage was sufficient to partially restore full-length receptor levels, as a significant increase in full-length p75^NTR^ (P = 0.0496) was detected in cells co-treated with dynasore compared to cells exposed to 6-OHDA alone (Fig. 2E). These findings indicate that receptor internalization is required for proteolytic processing of p75^NTR^ induced by oxidative stress.

**Figure 2.**
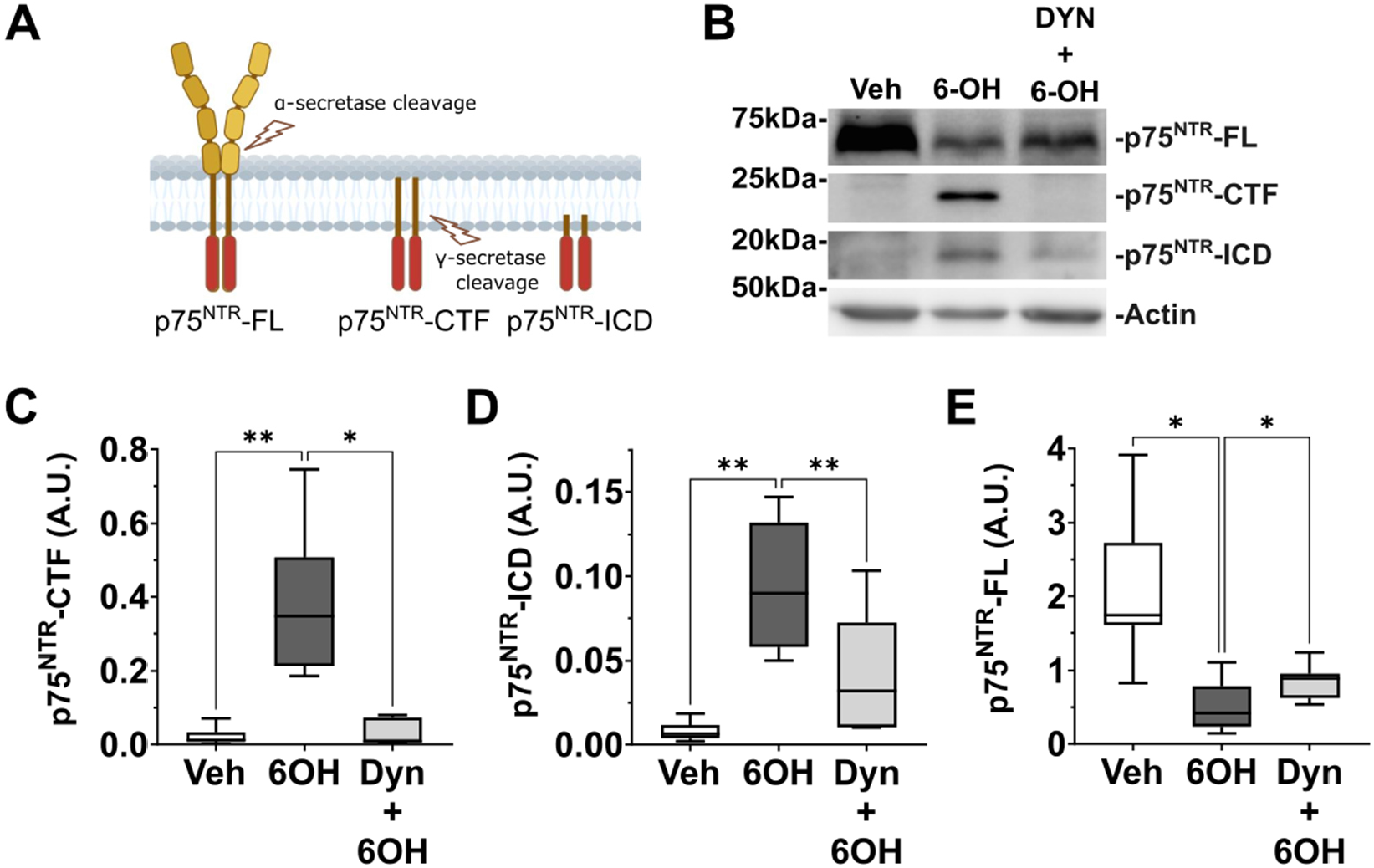
Receptor Internalization is Required for Cleavage of p75^NTR^ in Response to Oxidative Stress. **(A)** Schematic depicting regulated intramembrane proteolysis of p75^NTR^. The extracellular domain of the receptor is initially cleaved by an α-secretase, thereby yielding a p75^NTR^-CTF of ∼24 kDa. Cleavage of the transmembrane region of the p75^NTR^-CTF yields a p75^NTR^-ICD fragment of ∼ 19kDa. **(B)** Representative immunoblot analysis of p75^NTR^ fragments in lysates of differentiated LUHMES cells that were exposed for 18h to vehicle solution or 10 µM 6-OHDA and cotreated with vehicle solution or 80 µM dynasore. Immunoblotting was performed using an antibody specific for the p75^NTR^-ICD. Cropped regions indicate different exposure times. Immunoblotting for actin was performed as a loading control. **(C – E)** Densitometric analysis of p75^NTR^-CTF **(C)**, p75^NTR^-ICD **(D)**, and p75^NTR^-FL **(E)** from immunoblots described in 2B (*n* = 7; c, Friedman test with Dunn’s post-hoc analysis; d and e, ANOVA with Tukey’s HSD). Abbreviations: *Veh*, vehicle; *6OH*, 6-hydroxydopamine; *Dyn*, dynasore; *A.U*., arbitrary units; *, p<0.05; **, p<0.01; *n*, number of experiments, each featuring an independent cell culture preparation.

### Role of p75^NTR^ Coreceptors in Oxidative Stress-induced p75^NTR^ Processing

The p75^NTR^ can mediate diverse physiological effects through interactions with different coreceptors, including Trk receptors and sortilin. To investigate the potential involvement of such coreceptors in oxidative stress-induced p75^NTR^ signaling, we first analyzed the production of Trk receptors and sortilin in differentiated LUHMES cells, evaluating whether levels of such proteins are altered in response to oxidative stress. Western blot assays revealed that healthy, differentiated LUHMES cells express sortilin, yet sortilin levels are unaltered in cells exposed to 6-OHDA (Fig. 3A). Analyses of Trk receptors revealed that differentiated LUHMES cells express TrkA (Fig. 3B and 3C) but lack expression of TrkB and TrkC (Fig. 3D and 3E). Interestingly, in a dose-dependent manner, TrkA levels were significantly decreased in cells exposed to 6-OHDA compared to healthy cells treated with vehicle solution (5 µM 6-OHDA, P= 0.0001; 7.5 µM 6-OHDA, P< 0.0001), suggesting that TrkA signaling is reduced in response to oxidative stress (Fig. 3C).

**Figure 3.**
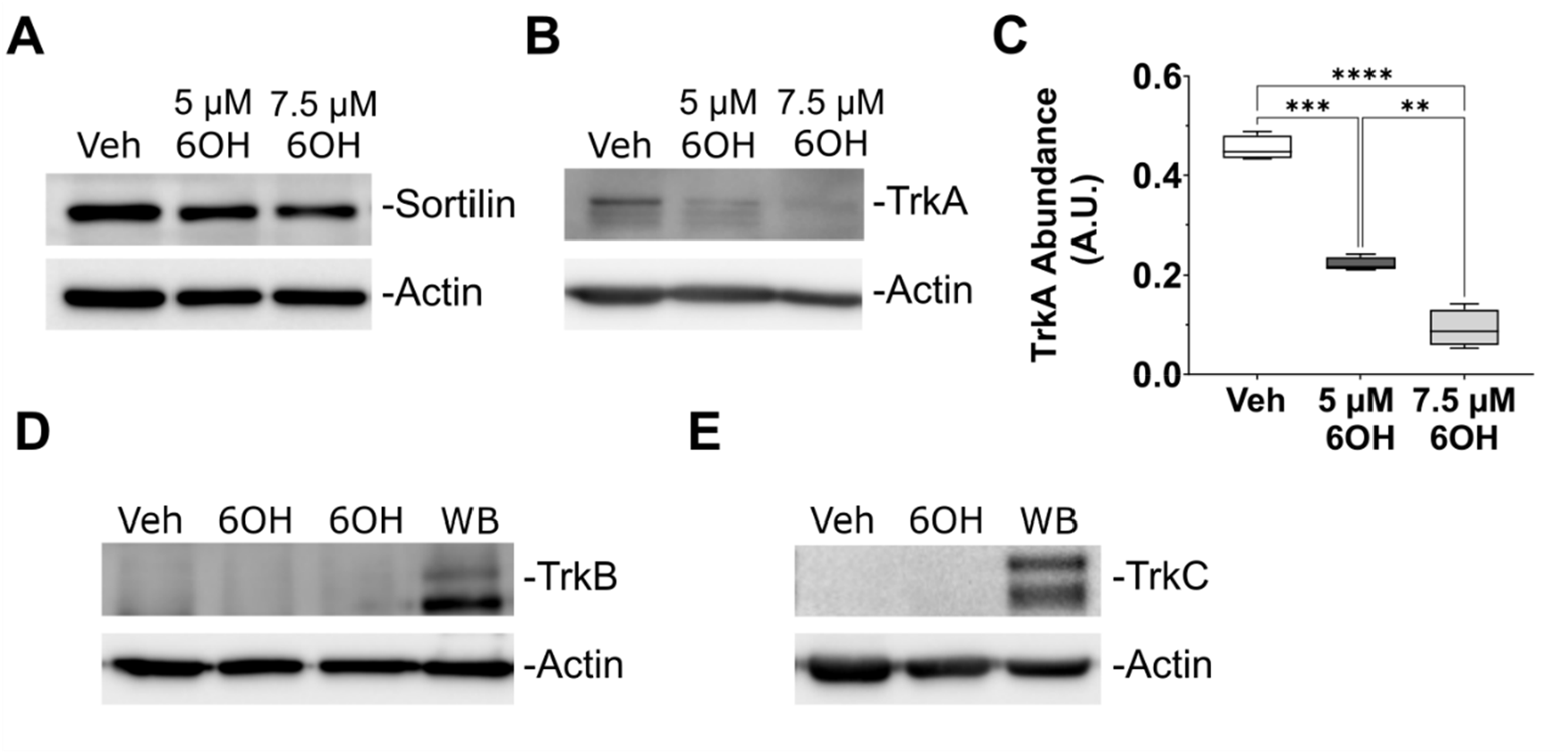
Effects of 6-Expression of p75^NTR^ Coreceptors in Dopaminergic Cells. **(A)** Representative immunoblot analysis of sortilin in lysates of differentiated LUHMES cells that were treated with vehicle solution, 5 µM 6-OHDA, or 7.5 µM 6-OHDA for 18h. Immunoblotting for actin was performed as a loading control. **(B)** Representative immunoblot analysis of TrkA in lysates of differentiated LUHMES cells that were treated with vehicle solution or 6-OHDA at the indicated concentrations for 18h. Immunoblotting for actin was performed as a loading control. **(C)** Densitometric analysis of TrkA from immunoblots as described in 3B (*n* = 4, ANOVA with Tukey’s HSD). **(D – E)** Representative western blot analysis of TrkB (D, *n =* 5) and TrkC (E, *n =* 3) in lysates of differentiated LUHMES cells exposed to vehicle solution or 5 µM 6-OHDA for 18h. Whole brain lysate was used as a positive control for Trk receptor detection, and immunoblotting for actin was performed as a loading control. Abbreviations: *Veh*, vehicle; *6OH*, 6-hydroxydopamine; *WB*, whole brain lysate; *A.U*., arbitrary units; **, p<0.01; ***, p<0.001; ****, p < 0.0001; *n*, number of experiments, each featuring an independent cell culture preparation.

**Figure 4.**
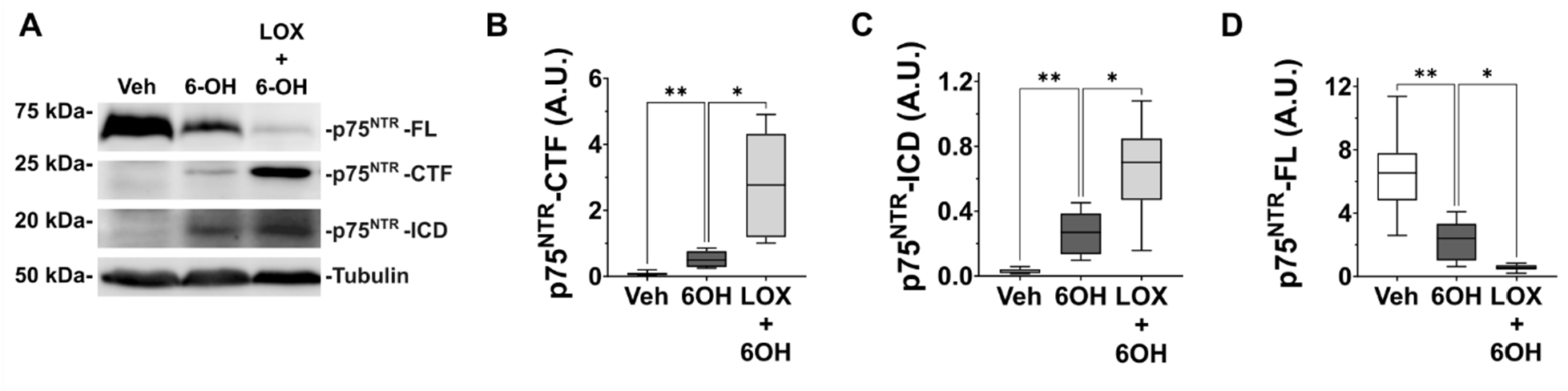
Trk Signaling Antagonizes p75^NTR^ Processing Induced by 6-OHDA. **(A)** Representative immunoblot of p75^NTR^ fragments from lysates of differentiated LUHMES cells treated with vehicle solution or 10 µM 6-OHDA following 3-hour pretreatment with vehicle solution or 5 µM LOXO-101. Immunoblotting was performed using an antibody specific for the p75^NTR^-ICD. Cropped regions indicate different exposure times. Immunoblotting for tubulin was performed as a loading control. (B – D) Densitometric analysis of p75^NTR^-CTF (B), p75^NTR^-ICD (C), and p75^NTR^-FL (D) from immunoblots described in 4A (*n* = 8; ANOVA with Tukey’s HSD). Abbreviations: *Veh*, vehicle; *6OH*, 6-hydroxydopamine; *LOX*, LOXO-101; *A.U*., arbitrary units; *, p<0.05; **, p<0.01; *n*, number of experiments, each featuring an independent cell culture preparation.

Based on our observation that TrkA levels decline in dopaminergic cells exposed to 6-OHDA, we next sought to evaluate whether reduced Trk signaling influences oxidative stress-induced p75^NTR^ processing. We therefore analyzed p75^NTR^ cleavage induced by 6-

### The Small Molecule p75^NTR^ Modulator, LM11a-31, Blocks Proteolytic Processing of p75^NTR^

We next sought to understand the physiological role of this signaling pathway and whether the cascade can be pharmacologically targeted to reduce neurodegeneration in a cell culture model of PD. LM11a-31, a small molecule ligand for p75^NTR^, has been reported to modulate p75^NTR^ signaling, but its mechanism of action remains incompletely understood. Thus, we explored the effects of LM11a-31 on p75^NTR^ cleavage induced by oxidative stress. Western blot analysis of p75^NTR^ fragments was performed on lysates of differentiated LUHMES cells that were exposed to 6-OHDA and cotreated with vehicle solution or LM11a-31. As expected, in cells exposed to 6-

Interestingly, despite the substantial decrease in p75^NTR^ fragments associated with LM11a-31 treatment, this reduction in p75^NTR^ cleavage was insufficient to restore full-length levels of the receptor, as we only detected a moderate, non-significant increase (P > 0.05) in p75^NTR^-FL in cells cotreated with LM11a-31 compared to cells treated with 6-OHDA alone (Fig. 5A and 5D).

**Figure 5.**
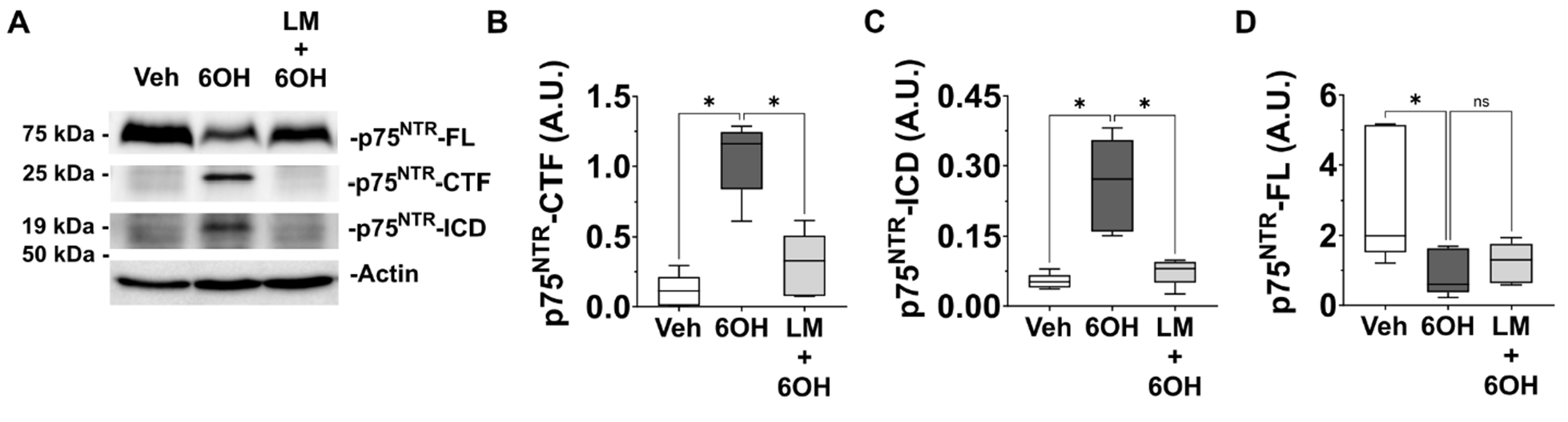
LM11a-31, a Small Molecule Modulator of p75^NTR^, Inhibits Proteolytic Processing of p75^NTR^ Induced by 6-OHDA. **(A)** Representative immunoblot of p75^NTR^ fragments from lysates of differentiated LUHMES cells treated with vehicle solution or 10 µM 6-OHDA and cotreated with vehicle solution or 20 nM LM11a-31. Immunoblotting was performed using an antibody specific for the p75^NTR^-ICD. Cropped regions indicate different exposure times. Immunoblotting for actin was performed as a loading control. (B – D) Densitometric analysis of p75^NTR^-CTF (B), p75^NTR^-ICD (C), and p75^NTR^-FL (D) from immunoblots described in 5A (*n* = 5; ANOVA with Tukey’s HSD). Abbreviations: *Veh*, vehicle; *6OH*, 6-hydroxydopamine; *LM*, LM11a-31; *A.U*., arbitrary units; *, p<0.05; *ns*, not significant; *n*, number of experiments, each featuring an independent cell culture preparation.

### LM11a-31 Protects Dopaminergic Cells from Neurodegeneration Associated with Oxidative Stress

In specific regions of the nervous system, p75^NTR^ exerts cell type-specific effects on neuronal survival, promoting death in certain neuronal populations yet enhancing survival in others[15, 33]. Thus, we next explored the physiological role of oxidative stress-induced p75^NTR^ signaling in dopaminergic cells derived from the ventral mesencephalon. Differentiated LUHMES cells were established in 8-well chamber slides. The cultures were treated with vehicle solution or 6-OHDA, and co-treatment was performed with vehicle solution or LM11a-31. Neurodegeneration induced by 6-OHDA was then assessed by nuclear labeling, fluorescence microscopy, and blinded scoring of pyknotic nuclei. Compared to healthy cells treated with vehicle solution, cultures exposed to 6-exhibited a significant increase in neuronal death (P < 0.0001). However, in cultures co-treated with LM11a-31, death of cells induced by 6-OHDA was significantly reduced (P = 0.0181) (Fig. 6A – B). Considered with our finding that LM11a-31 inhibits 6-OHDA-induced p75^NTR^ cleavage, these results suggest that p75^NTR^ cleavage promotes neuronal death associated with oxidative stress in dopaminergic cells. Moreover, our results collectively support a model in which oxidative stress stimulates JNK-mediated internalization of p75^NTR^ from the cell surface, thereby leading to cleavage of p75^NTR^ by endosomal proteases, while concurrently oxidative stress triggers the downregulation of Trk receptors that would otherwise antagonize p75^NTR^ processing.

**Figure 6.**
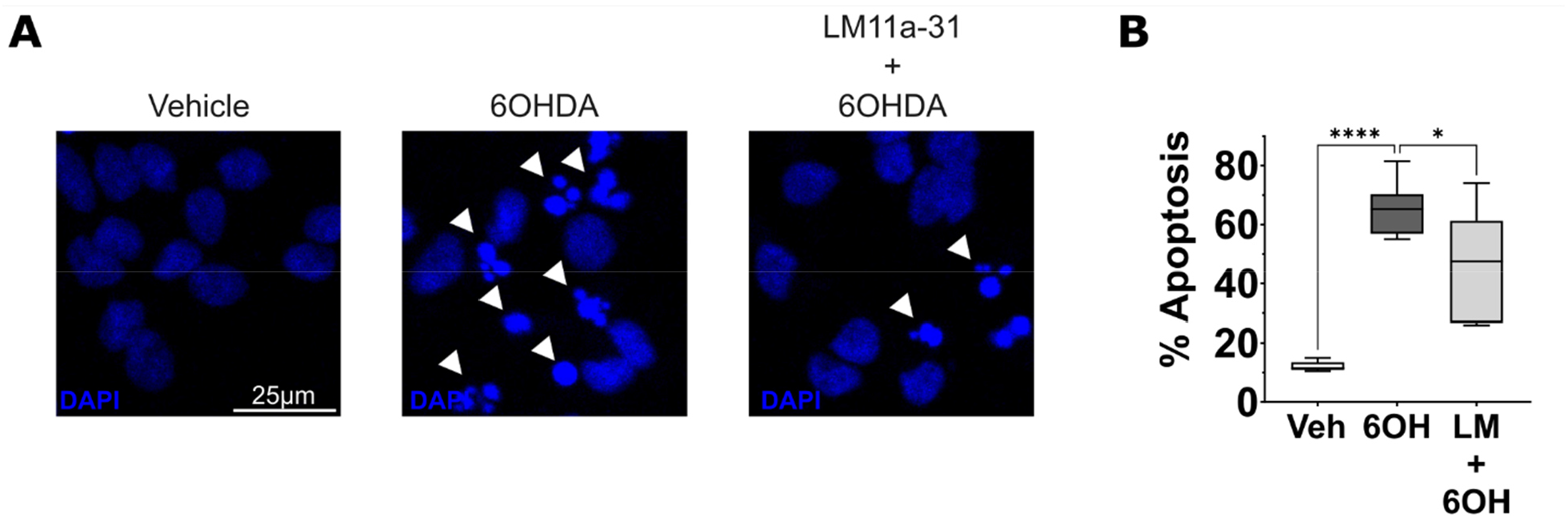
LM11a-31 Protects Dopaminergic Cells from Death Induced by 6-OHDA. **(A)** Confocal micrographs of differentiated LUHMES cells that were treated with vehicle solution or 7.5 µM 6-OHDA and cotreated with vehicle solution or 20 nM LM11a-31. Cultures were fixed 24H after treatment, and nuclei were labeled with the nucleic acid dye 4’,6-diamidino-2-phenylindole (DAPI) at a concentration of 5 µg/mL. Arrowheads indicate apoptotic cells with chromatin condensation, reduced nuclear area, and nuclear fragmentation. **(B)** Mean apoptosis of differentiated LUHMES cells that were treated and stained as described in 6A (*n* = 6; ANOVA with Tukey’s HSD). Abbreviations: *Veh*, vehicle; *6OH*, 6-hydroxydopamine; *LM*, LM11a-31; *, p<0.05; ****, p < 0.0001; *n*, number of experiments, each featuring an independent cell culture preparation.

These events lead to enhanced generation of p75^NTR^ fragments and induction of downstream pro-death signaling (Fig. 7).

**Figure 7.**
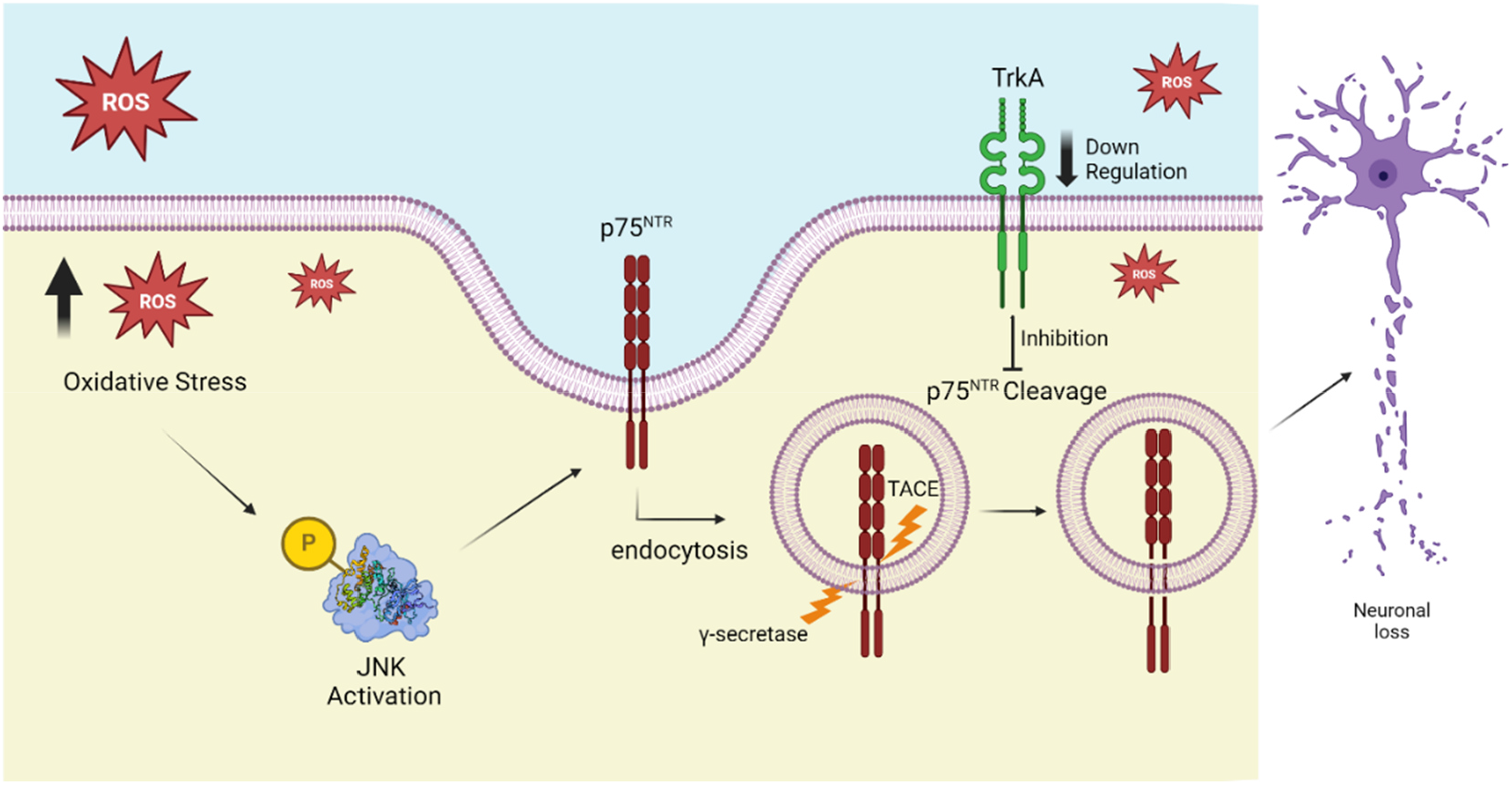
Schematic of Oxidative Stress-induced p75^NTR^ Signaling in Dopaminergic Cells. The accumulation of reactive oxygen species (ROS) and oxidative damage in cells with oxidative stress triggers activation of JNK. This activation stimulates clathrin-mediated endocytosis of p75^NTR^, increasing its susceptibility to cleavage by metalloproteases, such as TACE, and the γ-secretase complex in endosomes. Simultaneously, oxidative stress causes downregulation of Trk receptors, diminishing their antagonism of p75^NTR^ cleavage. Together, these processes result in accumulation of p75^NTR^ fragments, which contribute to downstream signaling pathways that promote neuronal death.

## Discussion

Neurodegeneration induced by p75^NTR^ has been linked to several pathological conditions associated with oxidative stress, including Alzheimer’s disease, Huntington’s disease, stroke, and amyotrophic lateral sclerosis[26, 29, 48-51]. Despite these associations, the signaling mechanisms underlying the effects of p75^NTR^ remain incompletely understood. Moreover, while p75^NTR^ is produced in injured dopaminergic neurons of the mesencephalon[35-38], a population vulnerable to oxidative stress and neurodegeneration in individuals with PD[52], the role of the receptor in PD remains unclear. In the present study, we identify JNK-mediated p75^NTR^ endocytosis and downregulation of Trk signaling as key steps in oxidative stress-induced p75^NTR^ signaling.

Additionally, using a cell culture model of PD, we reveal a pro-death role for p75^NTR^ and demonstrate that the receptor can be pharmacologically targeted to reduce neurodegeneration. These findings provide new insights into the role of p75^NTR^ in neurodegenerative diseases and highlight p75^NTR^ as a potential therapeutic target for mitigating neurodegeneration in PD.

Internalization of p75^NTR^ in response to neurotrophin binding is a key event in a variety of physiological contexts. For example, neurotrophin-induced internalization of p75^NTR^ regulates glioma progression[45], as well as death of sympathetic neurons during early neurodevelopment[42]. However, functional roles for cell-surface-localized p75^NTR^ have also been reported, including promotion of growth cone collapse in hippocampal neurons[44]. We previously reported that oxidative stress stimulates p75^NTR^ processing through a ligand-independent mechanism[39, 41], yet it has remained unclear whether ligand-independent p75^NTR^ signaling occurs at the cell surface or requires receptor internalization. Moreover, the relationship between receptor localization and p75^NTR^ signaling in neurons derived from the ventral midbrain has been poorly understood. Here, we demonstrate that receptor internalization is required for p75^NTR^ signaling in a cell culture model of PD, as pharmacological blockade of endocytosis resulted in decreased 6-OHDA-induced p75^NTR^ cleavage. These results highlight receptor endocytosis as a key mechanism governing ligand-independent p75^NTR^ signaling and an important process mediating p75^NTR^ signaling in dopaminergic cells.

We previously demonstrated that JNK is required for p75^NTR^ processing in dopaminergic cells subjected to oxidative stress[39]. However, the process through which JNK facilitates p75^NTR^ cleavage remained unclear. Here, we demonstrate that JNK stimulates endocytosis of p75^NTR^ from the cell surface, an event required for oxidative stress-induced p75^NTR^ processing. In sympathetic neurons, JNK-mediated internalization of p75^NTR^ has been demonstrated in response to BDNF treatment[42]. Our present findings suggest that JNK serves a similar role in response to pathological conditions associated with oxidative stress. Activation of JNK in response to oxidative stress has been reported in a variety of physiological contexts[53, 54], and increased JNK signaling has been found in models of other neurodegenerative diseases associated with oxidative stress, including Alzheimer’s disease[55] and amyotrophic lateral sclerosis[56]. Thus, our findings highlight the need for further studies evaluating whether JNK-induced endocytosis and cleavage of p75^NTR^ contributes to neurodegeneration in disorders associated with oxidative stress beyond PD.

Our studies utilized differentiated LUHMES cells subjected to oxidative stress as a model of midbrain dopaminergic neurons affected by PD. We characterized the expression of p75^NTR^ coreceptors in this model to determine how such coreceptors may influence oxidative stress-induced p75^NTR^ signaling. Our detection of TrkA in differentiated LUHMES cells aligns with multiple reports indicating that dopaminergic neurons in the ventral midbrain express TrkA or are responsive to the TrkA ligand NGF[57-59]. Importantly, we found that 6-OHDA induces downregulation of TrkA, indicating that TrkA signaling in dopaminergic cells is negatively regulated by oxidative stress.

Moreover, we discovered that downregulation of Trk signaling facilitates p75^NTR^ cleavage, as Trk receptor antagonism exacerbated oxidative stress-induced p75^NTR^ processing. Interestingly, TrkB expression has been reported in the ventral midbrain, including in dopaminergic neurons of the substantia nigra[60]. Thus, we were surprised that TrkB was undetectable in differentiated LUHMES cells, and such results may reveal a limitation of the LUHMES cell model. However, since TrkA and TrkB receptors are tyrosine kinase receptors with common downstream targets in the PI3K-Akt, ERK, and PLC cascades[14], our results suggest that in cells expressing TrkB, Trk signaling may similarly antagonize p75^NTR^ processing induced by oxidative stress. Further studies are needed to explore this hypothesis.

It is well-established that p75^NTR^ exerts cell type-specific effects, promoting survival in certain types of neurons while inducing death in others[14, 33]. Our results indicate that Trk signaling antagonizes p75^NTR^ processing in dopaminergic cells derived from the ventral mesencephalon, thereby limiting pro-death p75^NTR^ signaling. In contrast, Trk signaling was reported to potentiate p75^NTR^ cleavage in cerebellar granular neurons and PC12 cells, cell populations in which p75^NTR^ signaling is not pro-apoptotic[61, 62]. These observations suggest that Trk signaling may antagonize p75^NTR^ signaling to prevent neuronal death in certain cell populations, yet enhance p75^NTR^ signaling in other populations where p75^NTR^ signaling increases survival or differentiation. Further studies are needed to understand the molecular mechanisms governing the effects of Trk signaling on p75^NTR^ processing between cell types.

Although p75^NTR^ promotes proapoptotic signaling in response to activation by proneurotrophins, in certain physiological contexts the receptor may engage in ligand-independent signaling [21, 39, 41, 63-65]. Thus, treatment strategies for neurodegenerative conditions that involve inhibition of p75^NTR^-mediated apoptotic signaling should ideally mitigate both ligand-induced and ligand-independent activation of p75^NTR^. Here, we demonstrate that LM11a-31 decreases pro-death p75^NTR^ signaling in dopaminergic cells by reducing oxidative stress-induced p75^NTR^ cleavage. Interestingly, our previous studies demonstrated that oxidative stress stimulates p75^NTR^ cleavage through ligand-independent mechanisms[39, 41], and thus, the present findings highlight a novel function for LM11a-31 in inhibiting ligand-independent p75^NTR^ signaling. LM11a-31 is a small molecule based on the structural and chemical properties of a β-hairpin loop of NGF known to interact with p75^NTR^ [66]. Despite well-established, pro-survival effects of the compound in cells of the forebrain and spinal cord[67-69], its specific mechanisms of action are complex and incompletely understood. For example, LM11a-31 has been demonstrated to block proneurotrophin binding to p75^NTR^ and antagonize proneurotrophin-induced cell death in cultured oligodendrocytes[66], as well as in an *in vivo* model of spinal cord injury[68]; yet, multiple studies have supported that LM11a-31 is a biased agonist which functions in a manner distinct from NGF to shift p75^NTR^ interactions to stimulate pro-survival signaling while diminishing pro-death pathways[70-72]. Our current results suggest that, for cell types in which p75^NTR^ is pro-apoptotic, LM11a-31 can diminish ligand-independent p75^NTR^ signaling by decreasing the susceptibility of p75^NTR^ to enzymatic proteolysis. Further studies are needed to understand the specific interactions through which LM11a-31 modulates p75^NTR^ processing.

Although expression of p75^NTR^ in injured dopaminergic neurons of the ventral mesencephalon has been reported by several groups[35-38], the physiological impacts of the receptor in this neuronal population, and associated effects on PD, are incompletely understood. Pro-apoptotic signaling mediated by p75^NTR^ was observed in a study with primary cultures of dopaminergic neurons from mice with deficient expression of the pro-survival transcription factors Engrailed-1 and Engrailed-2[36]. However, environmental factors are thought to have a central role in PD onset and progression, since an estimated 90% of PD cases are sporadic[73], and thus further studies have been needed to understand whether p75^NTR^ contributes to dopaminergic neurodegeneration in PD models involving environmental factors that promote oxidative damage. Moreover, since oxidative damage can promote dopaminergic neurodegeneration through a multitude of pathways[52], it has remained unclear whether specifically modulating p75^NTR^ is sufficient to confer neuroprotection. Our present findings demonstrate that LM11a-31 prevents cleavage of p75^NTR^ and protects dopaminergic cells from neurodegeneration associated with oxidative stress. These results reveal p75^NTR^ as a key mediator of neuronal death induced by oxidative stress in dopaminergic cells and highlight the receptor as a potential pharmacological target for preventing dopaminergic neurodegeneration.

Multiple *in vivo* studies investigating the use of LM11a-31 to reduce neurodegeneration have yielded encouraging results. For example, *in vivo* administration of LM11a-31 was reported to reduce cholinergic neurite degeneration in a mouse model of Alzheimer’s disease[69], decrease hippocampal and cortical neuron death following traumatic brain injury[74], and enhance oligodendrocyte survival after spinal cord injury[68]. Moreover, a 26-week Phase 2 clinical trial was recently conducted to evaluate the safety of LM11a-31 as a treatment for Alzheimer’s disease. The study results indicated that the treatment was generally well tolerated, and exploratory outcome measures revealed significant Differences between drug- and placebo-treated groups, consistent with the hypothesis that the compound reduces AD progression[75]. Based on this accumulating evidence, studies evaluating potential therapeutic effects of the compound on other neuropathological conditions are needed. Our present findings reveal novel neuroprotective effects of LM11a-31 on dopaminergic cells derived from the ventral mesencephalon. The results highlight LM11a-31 as a potential PD treatment and underscore the need for *in vivo* studies evaluating the efficacy of LM11a-31 in preventing dopaminergic neurodegeneration.

## Acknowledgements

This research was supported by the National Institute of Neurological Disorders and Stroke (NINDS) [1R15NS111413-01A1], as well as by the University of Alabama in Huntsville Summer Community of Scholars (SCS) program. The authors also thank Dr. Bruce Carter for generously donating p75^NTR^-ICD antibody for these studies, as well as Isaac Perkins for contributions to image analyses.

## Author Contributions

P.V.K. performed experiments, analyzed data, and contributed to written portions of the manuscript. A.M.N. and S.C.O. performed experiments, analyzed data, and reviewed the manuscript. L.G.D. and C.A.S. performed experiments and reviewed the manuscript. B.R.K. designed the experiments, performed experiments, analyzed data, and drafted the article.

## Data Availability

Data that support the findings of this study are available from Dr. Bradley Kraemer, the corresponding author, upon reasonable request.

